# Expression Level Analysis of ACE2 Receptor Gene in African-American and Non-African-American COVID-19 Patients

**DOI:** 10.1101/2023.09.11.557129

**Authors:** Marion N. Nyamari, Kauthar M. Omar, Ayorinde F. Fayehun, Oumaima Dachi, Billiah Kemunto Bwana, Olaitan I. Awe

## Abstract

**Background:** The COVID-19 pandemic caused by SARS-CoV-2 has spread rapidly across the continents. While the incidence of COVID-19 has been reported to be higher among African-American individuals, the rate of mortality has been lower compared to that of non-African-Americans. ACE2 is involved in COVID-19 as SARS-CoV-2 uses the ACE2 enzyme to enter host cells. Although the difference in COVID-19 incidence can be explained by many factors such as low accessibility of health insurance among the African-American community, little is known about ACE2 expression in African-American COVID-19 patients compared to non-African-American COVID-19 patients. The variable expression of genes can contribute to this observed phenomenon.

**Methodology:** In this study, transcriptomes from African-American and non-African-American COVID-19 patients were retrieved from the sequence read archive and analyzed for ACE2 gene expression. HISAT2 was used to align the reads to the human reference genome, and HTseq-count was used to get raw gene counts. EdgeR was utilized for differential gene expression analysis, and enrichR was employed for gene enrichment analysis.

**Results:** The datasets included 14 and 33 transcriptome sequences from COVID-19 patients of African-American and non-African-American descent, respectively. There were 24,092 differentially expressed genes, with 7,718 upregulated (log fold change > 1 and FDR 0.05) and 16,374 downregulated (log fold change −1 and FDR 0.05). The ACE2 mRNA level was found to be considerably downregulated in the African-American cohort (p-value = 0.0242, p-adjusted value = 0.038).

**Conclusion:** The downregulation of ACE2 in the African-American cohort could indicate a correlation to the low COVID-19 severity observed among the African-American community.

## Introduction

Coronavirus Disease 2019 (COVID-19) is a viral ailment caused by Severe Acute Respiratory Syndrome Coronavirus 2 (SARS-CoV-2). Wuhan city, China was the first place where the virus was detected, before it spread to other countries throughout the world, causing a global pandemic [1]. Fever, dry cough, and malaise are common COVID-19 symptoms [1]. Difficulty breathing, chest pain as well as Acute Respiratory Distress Syndrome (ARDS) have been reported in severe cases [1].

As of August 1, 2020, the WHO recorded about 18 million COVID-19 cases and 600,000 deaths globally [2]. The total number of COVID-19 cases in the European region and United States were about 3 million and 4 million confirmed cases, out of a population of 741 million and 327 million respectively [3]. The rate of mortality among the African-American community was reported to be 25 percent lower than white and native Americans while Asian Americans had the lowest rates [4]. Areas with a large African-American population however, reported that COVID-19 patients had increased in-hospital mortality, caused by more comorbid conditions being present [5]. Older residents, people in dense housing such as the projects and African-Americans with heart disease and lacking health insurance had a doubled chance of dying from COVID-19 [6], [7]. Patient outcomes were improved by starting dialysis early and using ACE inhibitors prior to admission [5].

Asia with about 60% of the world population recorded about 28.6% of cases and 22.7% of deaths (303 deaths/million) as of 1^st^ of April 2022 [8]. According to the same report by worldometer, North America (4.7% of world population) appears to have been the worst hit with 19.7% and 23.4% of cases and deaths respectively (3,914 deaths/million). This is closely followed by Europe (9.6% of the world population) with 36.7% of cases and 28.7% of global cases and deaths respectively (2,374 deaths/million). Study reports have shown that next-generation sequences can be used to screen for disorders in newborns [9], for the study of pathogen evolution using African SARS-CoV-2 sequences [18], prostate cancer biomarker discovery [10], malaria/CoVID-19 biomarker discovery [58], SARS-CoV-2 variants classification [59], HIV-1 evolution in sub-Saharan Africa [60], analysis of RNA-seq and ChIP-seq data [61], and ebola virus comparative genomics [62]. We will be using next-generation sequencing data for our investigation in this study.

Zeberg and his colleagues showed that the major genetic risk to COVID-19 is caused by a 50-kilobase pair genomic segment inherited from the Neanderthals, which is present in 50% and 16% of people in south Asia and Europe respectively [12]. Gene allele distribution could also explain some of the differences in COVID-19 infection incidence among people in different geographic regions [3]. The authors however suggest that due to a lack of sufficient data, more genetic investigations are needed to corroborate this notion.

Angiotensin Converting Enzyme 2 (ACE2) hydrolyzes Ang-II to produce Ang-(1-7) - a vasodilator and anti-inflammatory agent, by. ACE2 promotes a protective anti-inflammatory response in the lung by lowering oedema, vascular permeability, and pulmonary cell damage. ACE2 promotes a protective anti-inflammatory response in the lung by lowering oedema, pulmonary cell damage and vascular permeability [16], [39], [50]. When the expression of ACE2 receptors is reduced, its substrate Ang-II, accumulates leading to increased vascular permeability and pulmonary oedema [3].

While ACE supports vasoconstriction and inflammation through Ang-II induction, ACE2 promotes vasodilation and anti-inflammatory responses through Ang-II breakdown. Thus, ACE and ACE2 play opposite roles in the generation and degradation of Ang-II and RAS tissue inflammatory balance, and ACE/ACE2 imbalance is believed to play a role in COVID-19 pathogenesis [17].

Genetics is believed to have a substantial control on plasma ACE levels [38]. There is a genetic deletion/insertion polymorphism of ACE enzyme, which is linked to half of the variation in the total plasma concentration of ACE [40]. The ACE I/D polymorphism of the Alu repeat in intron 16, consists of either an insertion (I) allele or a deletion (D) allele of *Alu* repeat, leading to three possible genotypes: II (homozygous insertion), ID (heterozygous - insertion of Alu repeat in one allele and deletion of the Alu repeat in the other allele) and DD (homozygous deletion) [40]. Individuals with D/D genotype were reported to have significantly higher serum ACE concentrations compared to the I/D and I/I genotypes [40]. The dominant DD allele is associated with elevated levels of serum and tissue ACE, and subsequently increase in Ang-II levels [3].

Delanghe and his colleagues showed that variation in the distribution of D/I genotype was associated with prevalence of COVID-19 in 25 different European countries, where countries with higher ACE D allele frequency had lower prevalence of COVID-19 [54]. Similarly, nations with high COVID-19 prevalence, such as China and Korea, which were both heavily affected by the virus in the outset, have low ACE D allele frequencies [43].

SARS-CoV-2 virus enters host cells via the ACE2 receptor [20], [3]. ACE2 receptor is a type I transmembrane amino-peptidase found on cell surfaces of the gut, heart, lungs, and many other organs,and also found in solution in blood plasma and urine [25], [3]. The COVID-19 viral envelope’s “spike” (S) protein binds to ACE2 receptors on nasopharyngeal mucosa and alveolar pneumocytes [20], [21], [51].

ACE2 expression is regulated by serum level of Ang-II through the AT1R, as Ang-II induces ACE in a dose dependent manner and decreases ACE2 expression [50]; Since ACE induces the production of Ang-II, high levels of Ang-II could therefore reduce ACE2 expression and activity.

Therefore, in ACE-DD homozygous individuals with increased ACE activities and Ang-II level, ACE2 expression might be down-regulated [17]. Furthermore, populations with this DD polymorphism may have reduced ACE2 expression which may play a protective role against COVID-19.

ACE2 genes contain 18 exons and maps on Xp22. Genetic variations in ACE2 gene have been proposed to also affect the COVID-19 viral spike protein interaction with the ACE2 receptor [31]. Several ACE2 gene variations can affect interaction with spike protein, protein stability, or change in ligand receptor affinity [19]. Therefore it can be inferred that genetic variations in the ACE2 gene may influence the individual’s susceptibility or resistance to SARS-CoV-2 [19], [23]. Search for variants that show a correlation with clinical COVID-19 severity may explain the broad range of SARS-CoV-2 infection outcome. COVID-19. CoVID-19 symptoms and outcomes might be associated with the pattern and level of human ACE2 enzyme expression in different tissues [53]. Also, circulating ACE2 might inhibit SARS-COV-2 entry into pulmonary alveoli and CoVID-19 infection severity correlating with soluble/membrane bound ratio of ACE2 [15].

ACE2 polymorphism, which reduces ACE2 receptor expression, is consequently expected to be associated with increased Ang-II levels. Population distribution of these alleles may also explain the distribution of CoVID-19. However, many other studies have not been able to find this association. For example, a study of three ACE2 polymorphisms (rs2106809, rs4646155, and rs879922) in hypertensive patients did not show any significant relationship with circulating levels of ACE2 [32]. Also, an Italian study did not find a strong association between COVID-19 disease severity and ACE2 variants [35]. Furthermore, the SARS-CoV-2 virus may have an impact on the expression of ACE2 receptors, where it is downregulated in infected cells [28]. Also, the function of the ACE2 is likely impaired during the initial SARS-CoV-2 virus binding from ACE2 steric domain hindrance or ACE2 downregulation [13].

In the light of the potential role of ACE2 and Ang-II in the SARS-CoV-2 pathogenesis the quantification of ACE2 and Ang-II should be part of the COVID-19 patients’ biological assessment and monitoring [13]. In this study, we compared the ACE2 gene expression level in patients of African-American descent with COVID-19, to that of non-African-American COVID-19 patients.

## Methods

### Data Acquisition

Transcriptomic data of both African-American and Non-African-American populations was downloaded from NCBI. The reads were downloaded via their accession numbers from the command line using Fasterq-dump (https://trace.ncbi.nlm.nih.gov/Traces/sra/sra.cgi?view=software). The non-African-American population was obtained from bioproject <u>PRJNA743862</u>; GEO: GSE179448 while the African-American project was obtained from bioproject <u>PRJNA679264</u>; GEO: GSE161731. The African-American dataset included patients with COVID-19 as well as seasonal COVID.

### Quality Control

FastQC version 0.11.3 was used to verify the quality of raw sequencing reads [46], and subsequently trimming to eliminate adapter sequences was done Cutadapt [33].

### Alignment to the human reference genome

After passing the quality check, the reads were aligned to the human reference genome (hg38) using Hisat2 [26]. Samtools v1.3.1 [30] was used to convert the aligned reads’ SAM file to a BAM file, which was then sorted.

### Read counts

HTSeq-count [37] was used to count reads associated with each human gene.

### Normalization

EdgeR [41] was used to normalize library sizes using Trimmed Mean of M-values (TMM) by determining the scaling factor to obtain effective library sizes. Counts were then adjusted by effective library sizes to create counts per million (cpm) values.

### Differential gene expression analysis

EdgeR [41] was used to perform differential expression analysis according to established principles. Genes having corrected P-values of 0.05 and absolute fold increases of 1.5 were considered differentially expressed. Differentially expressed candidate genes were evaluated between African-American and non-African-American populations. Enrichr was used to perform Gene Ontology (GO) and KEGG pathway enrichment analysis of differentially expressed genes [14]. The data analysis workflow is shown in Figure 1 below

**Figure 1:**
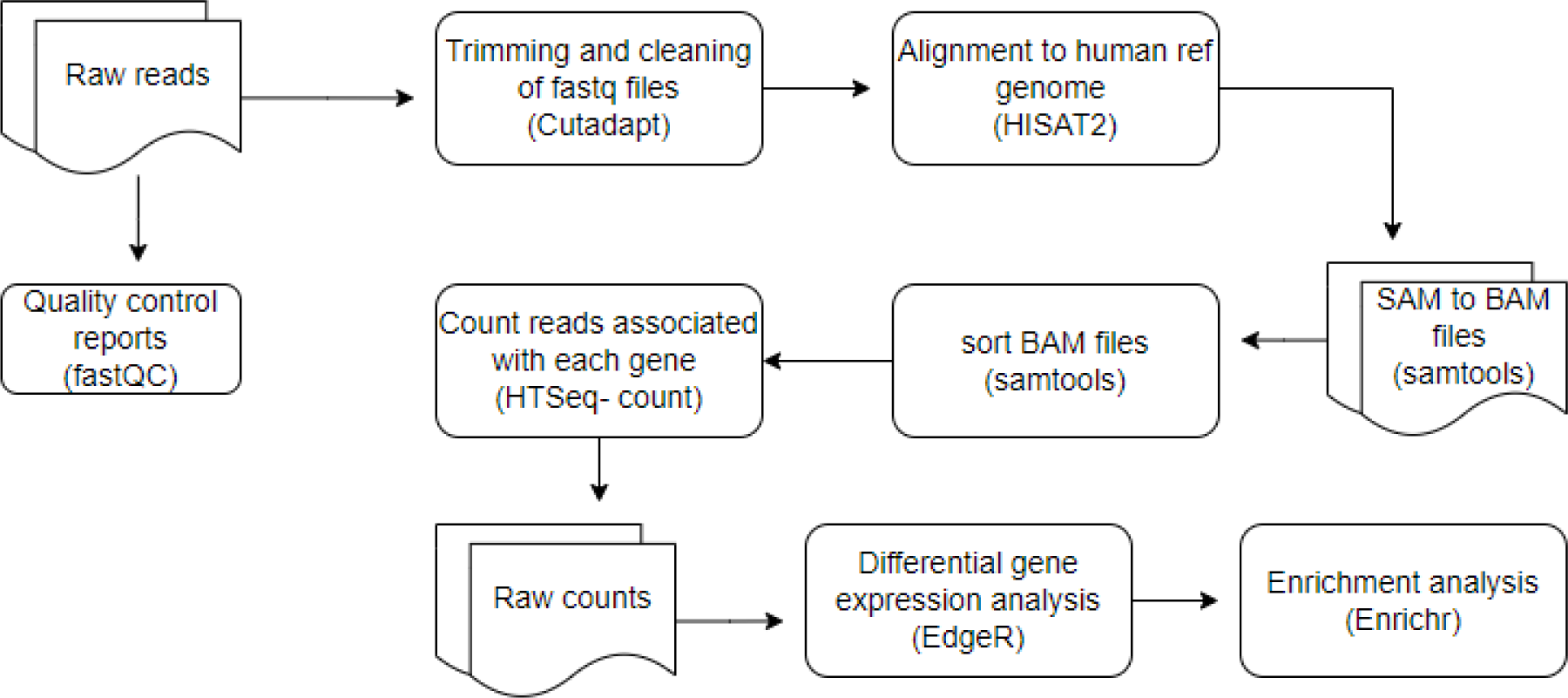
Materials and methods used in data analysis

## Results

### Quality control

The datasets included 14 and 33 transcriptomic sequences of African-American and non-African-American COVID-19 patients, respectively. The sequences were of high quality, all the sequences scored a Phred score of above 30 after trimming of the adaptor sequence.

### Identification of differentially expressed genes in the African-American population

The raw gene count matrix consisting of a total of 39,590 genes generated in the gene count step of our pipeline, was used in EdgeR to analyze differential expression levels of the genes. Differential expression analysis revealed 7,718 out of 24,092 genes were significantly upregulated (log fold change > 1 and FDR < 0.05) while 16,374 genes were significantly downregulated (log fold change < −1 and FDR < 0.05) between African-American and non-African-American populations (controls) (Figure.2 and 3). We discovered that the mRNA level of ACE2 was significantly downregulated in the African-American cohort. (p value = 0.0242, p-adjusted value = 0.038).

**Figure 2.**
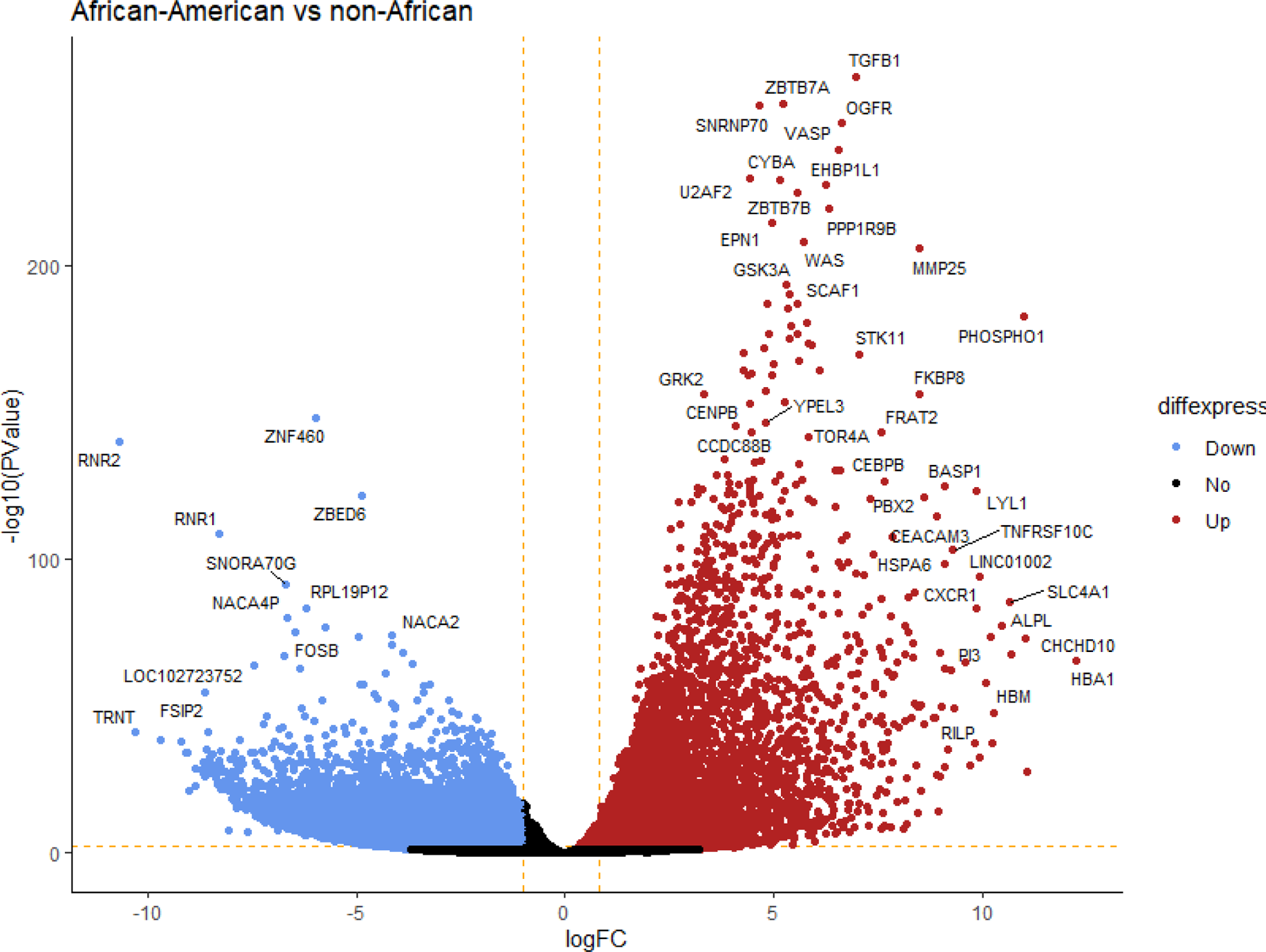
Volcano Plot for differentially expressed genes between African-American COVID-19 patients vs non-African-American COVID-19 patients as controls. Scattered dots represent genes; the x-axis is the log fold change for the ratio of African-American vs non-African-American COVID-19 patients, whereas the y-axis is the −log 10 (P Value) where P is the probability that a gene has statistical significance in its differential expression. Red dots represent genes significantly upregulated, blue dots are genes significantly suppressed genes while black dots are genes with non-significant changes between the conditions

**Figure 3.**
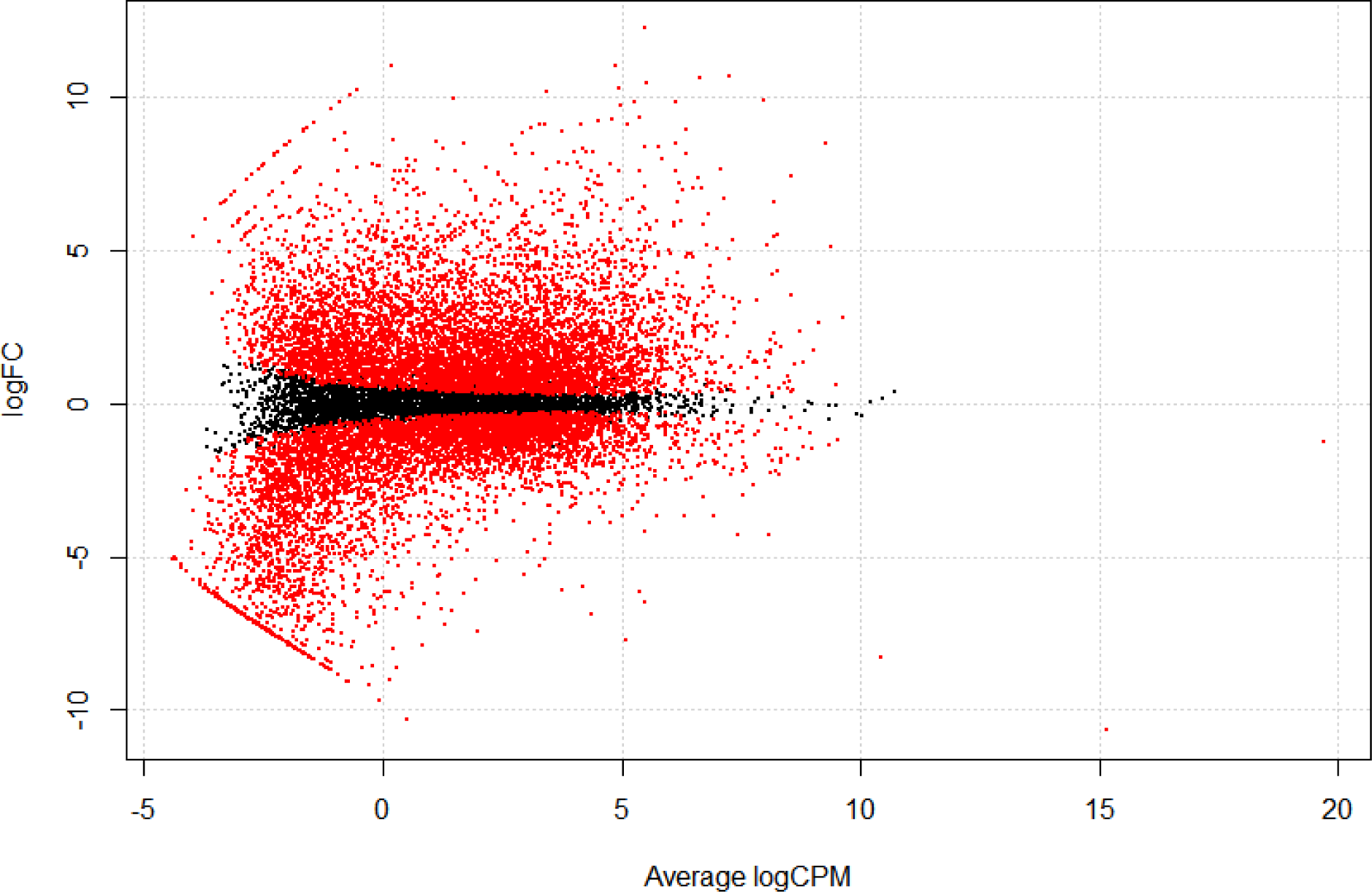
MA plot for transcript distribution determined by EdgeR. Red points represent differentially expressed transcripts (P-adjusted < 0.05).

The top 10 most significantly upregulated genes include TGFB1, ZBTB7A, SNRNP70, OGFR, VASP, U2AF2, CYBA, EHBP1L1, ZBTB7B, and PPP1R9B with a P-adjusted value < 0.05 (Table 1). The top 10 most significantly down regulated genes include ZNF460, RNR2, ZBED6, RNR1, SNORA70G, RPL19P12, NACA4P, RPL23P8, FOSB, NACA2 with a P-adjusted value < 0.05 (Table 2).

**Table. 1.** Top 10 significantly upregulated genes between African-American COVID-19 patients vs non-African-American COVID-19 patients.

| **Gene** | **Annotation\*** | **Log Fold Change** | **P-Value** | **FDR P-Value** |
| --- | --- | --- | --- | --- |
| TGFB1 | Transforming Growth Factor Beta 1 | 6.947 | 4.91E-265 | 1.83E-260 |
| ZBTB7A | Zinc Finger and BTB Domain Containing 7A | 5.201 | 6.88E-256 | 1.28E-251 |
| SNRNP70 | Small Nuclear Ribonucleoprotein U1 Subunit 70 | 4.643 | 6.98E-255 | 8.66E-251 |
| OGFR | Opioid Growth Factor Receptor | 6.609 | 5.98E-249 | 5.57E-245 |
| VASP | Vasodilator Stimulated Phosphoprotein | 6.547 | 5.25E-240 | 3.91E-236 |
| U2AF2 | U2 Small Nuclear RNA Auxiliary Factor 2 | 4.408 | 1.64E-230 | 1.02E-226 |
| CYBA | Cytochrome B-245 Alpha Chain | 5.159 | 5.82E-230 | 3.10E-226 |
| EHBP1L1 | EH Domain Binding Protein 1 Like 1 | 6.234 | 4.15E-228 | 1.93E-224 |
| ZBTB7B | Zinc Finger and BTB Domain Containing 7B | 5.573 | 1.44E-225 | 5.96E-222 |
| PPP1R9B | Protein Phosphatase 1 Regulatory Subunit 9B | 6.335 | 8.13E-220 | 3.03E-216 |

**Table. 2.**
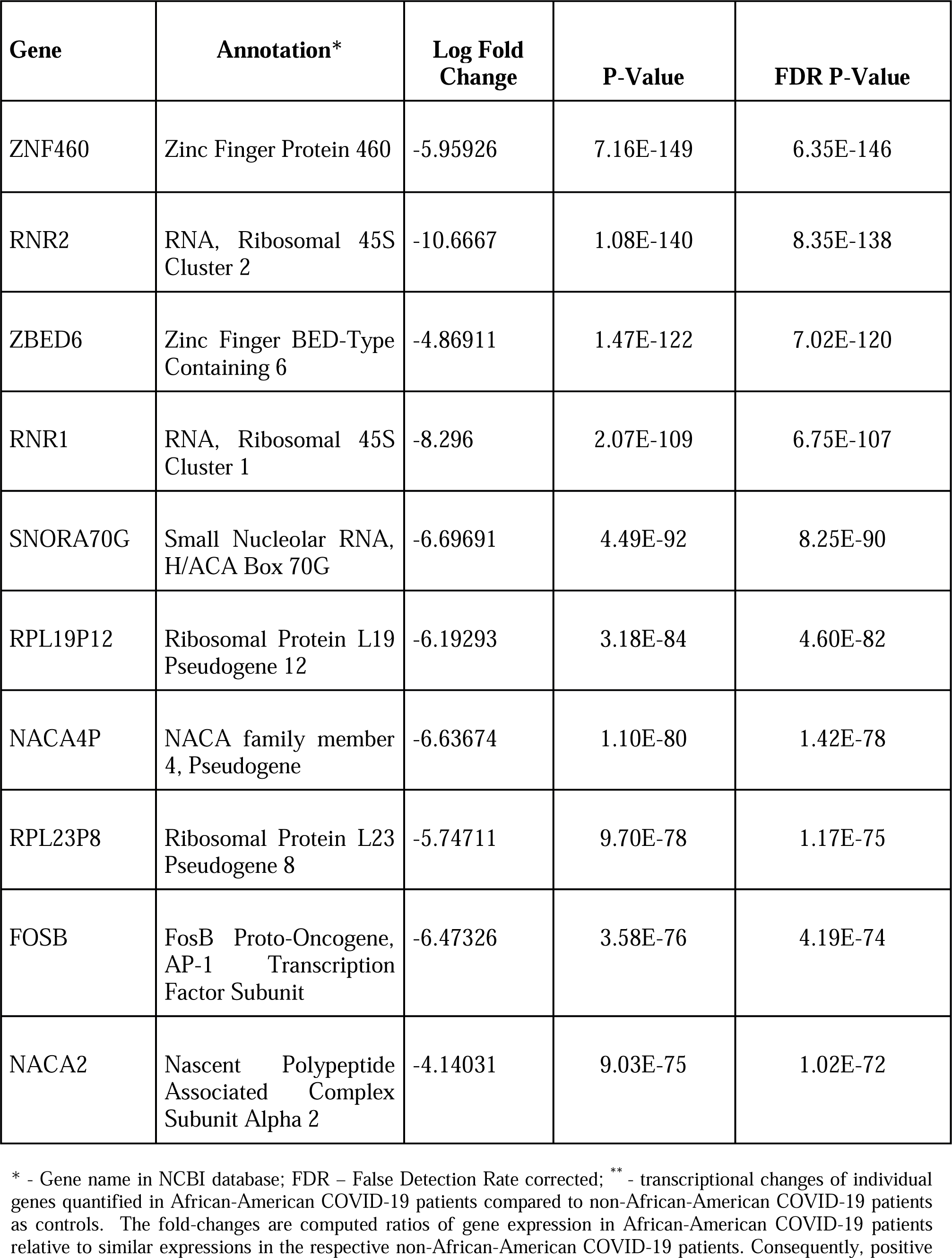
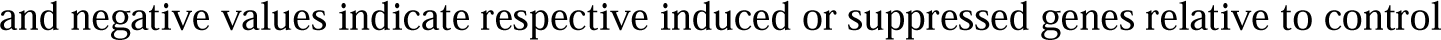
Top 10 significantly downregulated genes between African-American COVID-19 patients vs non-African-American COVID-19 patient.

| **Gene** | **Annotation\*** | **Log Fold Change** | **P-Value** | **FDR P-Value** |
| --- | --- | --- | --- | --- |
| ZNF460 | Zinc Finger Protein 460 | -5.95926 | 7.16E-149 | 6.35E-146 |
| RNR2 | RNA, Ribosomal 45S Cluster 2 | -10.6667 | 1.08E-140 | 8.35E-138 |
| ZBED6 | Zinc Finger BED-Type Containing 6 | -4.86911 | 1.47E-122 | 7.02E-120 |
| RNR1 | RNA, Ribosomal 45S Cluster 1 | -8.296 | 2.07E-109 | 6.75E-107 |
| SNORA70G | Small Nucleolar RNA, H/ACA Box 70G | -6.69691 | 4.49E-92 | 8.25E-90 |
| RPL19P12 | Ribosomal Protein L19 Pseudogene 12 | -6.19293 | 3.18E-84 | 4.60E-82 |
| NACA4P | NACA family member 4, Pseudogene | -6.63674 | 1.10E-80 | 1.42E-78 |
| RPL23P8 | Ribosomal Protein L23 Pseudogene 8 | -5.74711 | 9.70E-78 | 1.17E-75 |
| FOSB | FosB Proto-Oncogene, AP-1 Transcription Factor Subunit | -6.47326 | 3.58E-76 | 4.19E-74 |
| NACA2 | Nascent Polypeptide Associated Complex Subunit Alpha 2 | -4.14031 | 9.03E-75 | 1.02E-72 |

\* - Gene name in NCBI database; FDR – False Detection Rate corrected; \*\* - transcriptional changes of individual genes quantified in African-American COVID-19 patients compared to non-African-American COVID-19 patients as controls. The fold-changes are computed ratios of gene expression in African-American COVID-19 patients relative to similar expressions in the respective non-African-American COVID-19 patients. Consequently, positive
and negative values indicate respective induced or suppressed genes relative to control

An unsupervised hierarchical analysis revealed that sets of genes with comparable expression characteristics clustered together, as expected. Groups of genes are systematically, either up-regulated (red) or down-regulated (green) in African-Americans versus non-African-Americans (Figure 4).

**Figure 4.**
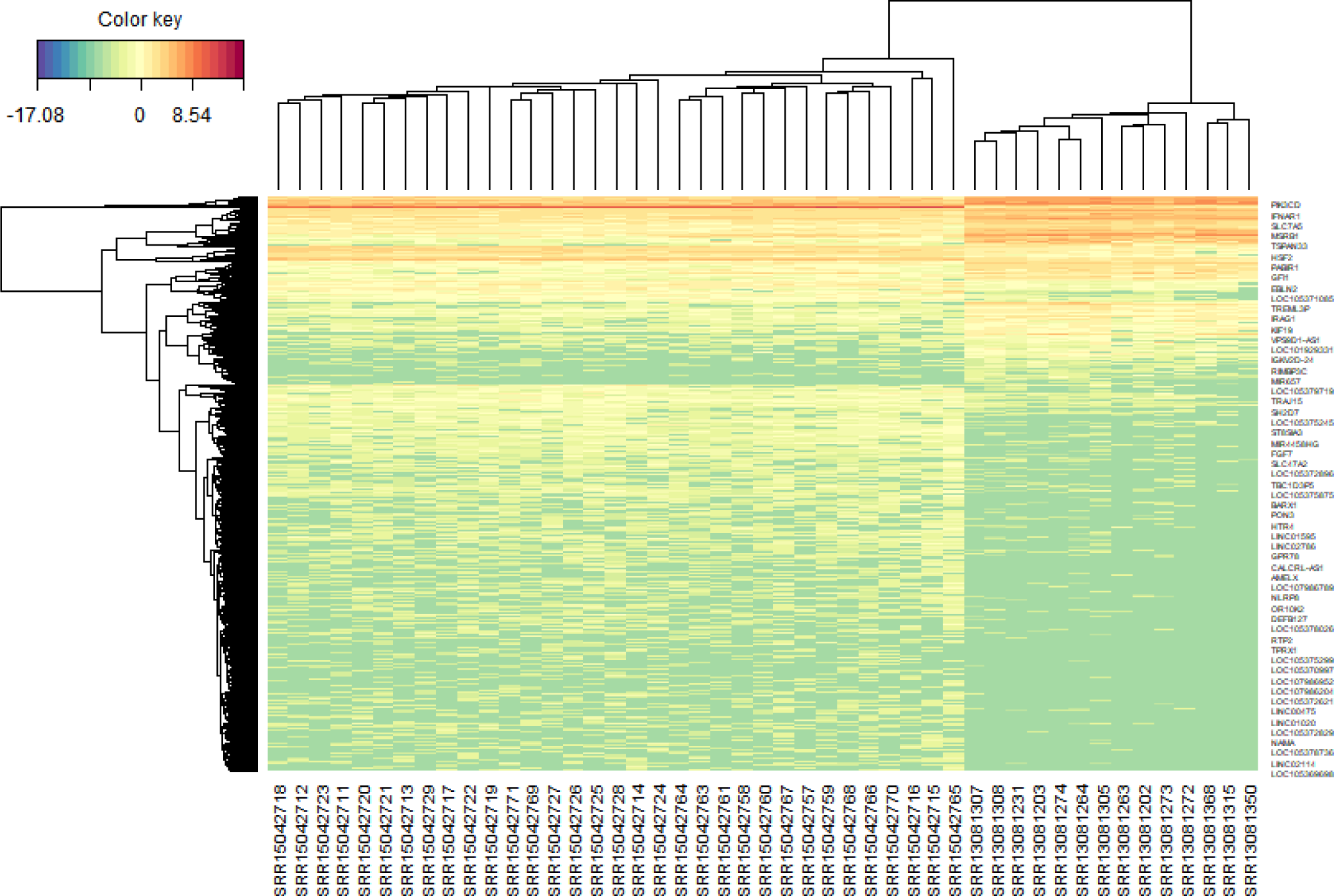
HeatMap for hierarchical clustering of differential gene expression between African-American COVID-19 patients vs non-African-American COVID-19 patients as controls. Groups of genes are systematically, either up-regulated (red) or down-regulated (green) in treatment versus controls.

### Functional analysis of significantly upregulated genes

Enrichment analysis was performed on significantly over-expressed genes to verify the molecular functions, biological processes, cellular components, and pathways associated with the differentially expressed genes. The fifteen most significant molecular functions, biological processes, cellular components, and ten most significant pathways associated with the differentially expressed genes are represented according to Gene Ontology (GO) biological process, Gene Ontology (GO) molecular function, Gene Ontology (GO) cellular component, and KEGG pathway analysis based on p value (Figure 5A-5D).

**Figure 5A.**
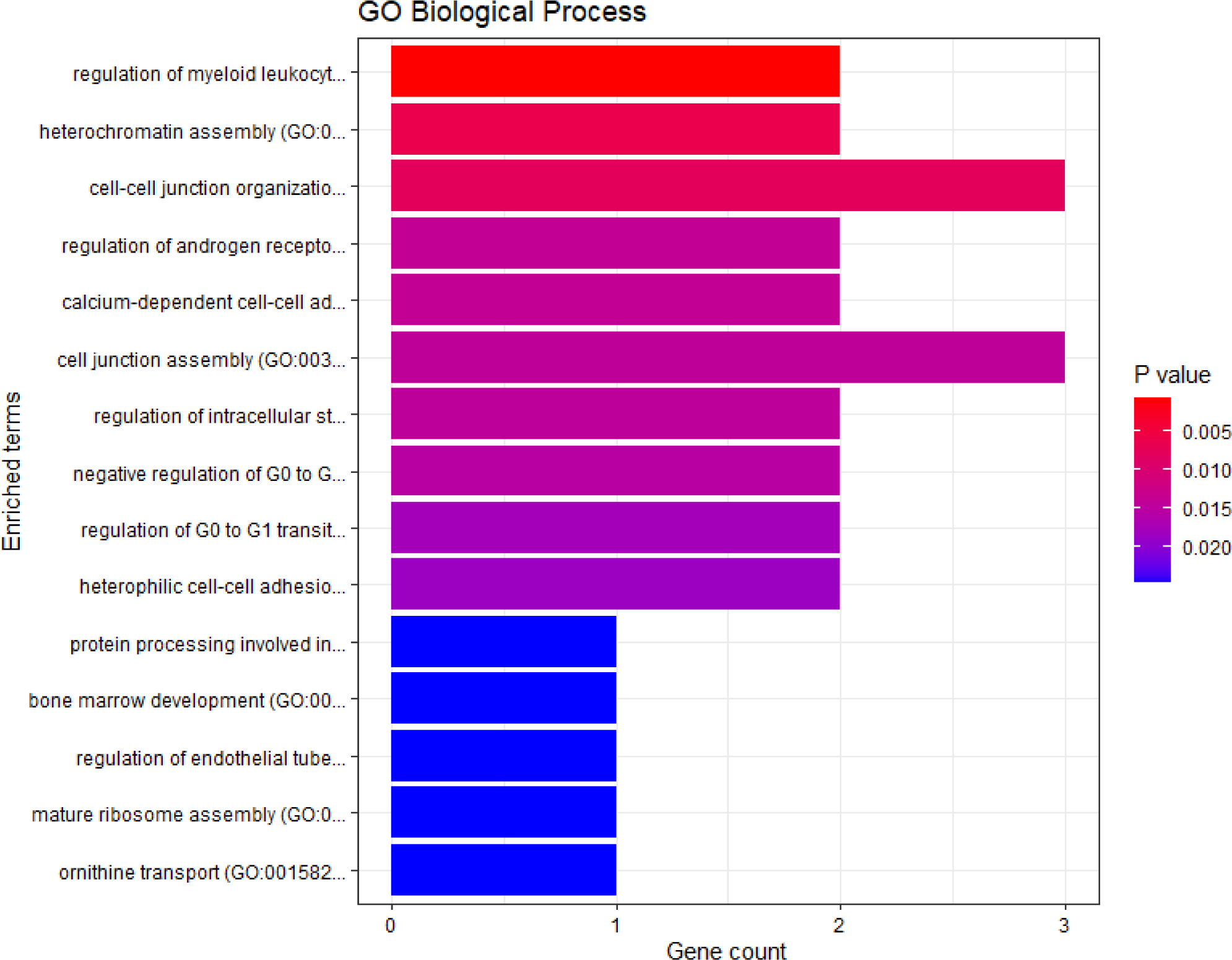
GO biological process

**Figure 5B.**
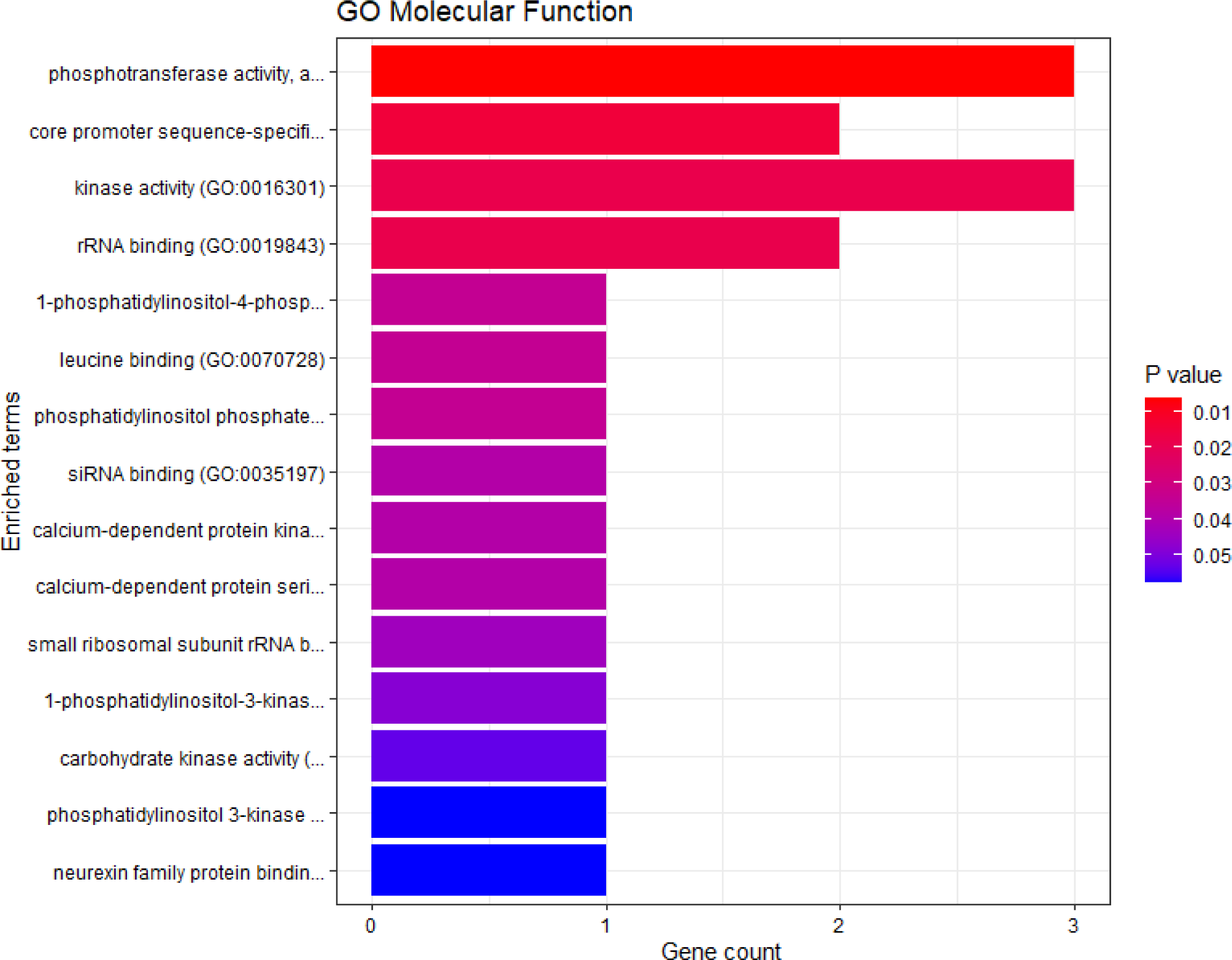
GO molecular function

**Figure 5C.**
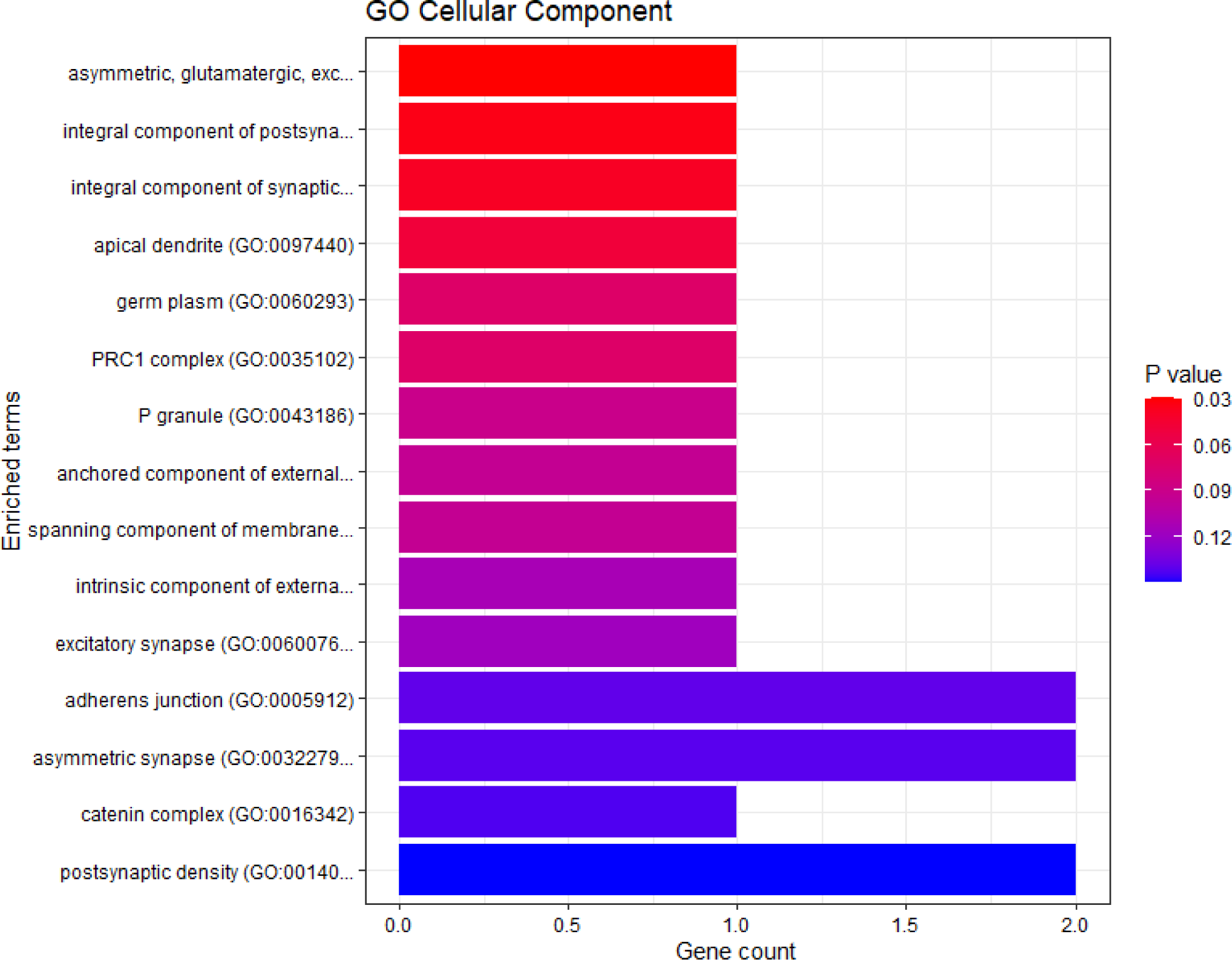
GO cellular component

**Figure 5D.**
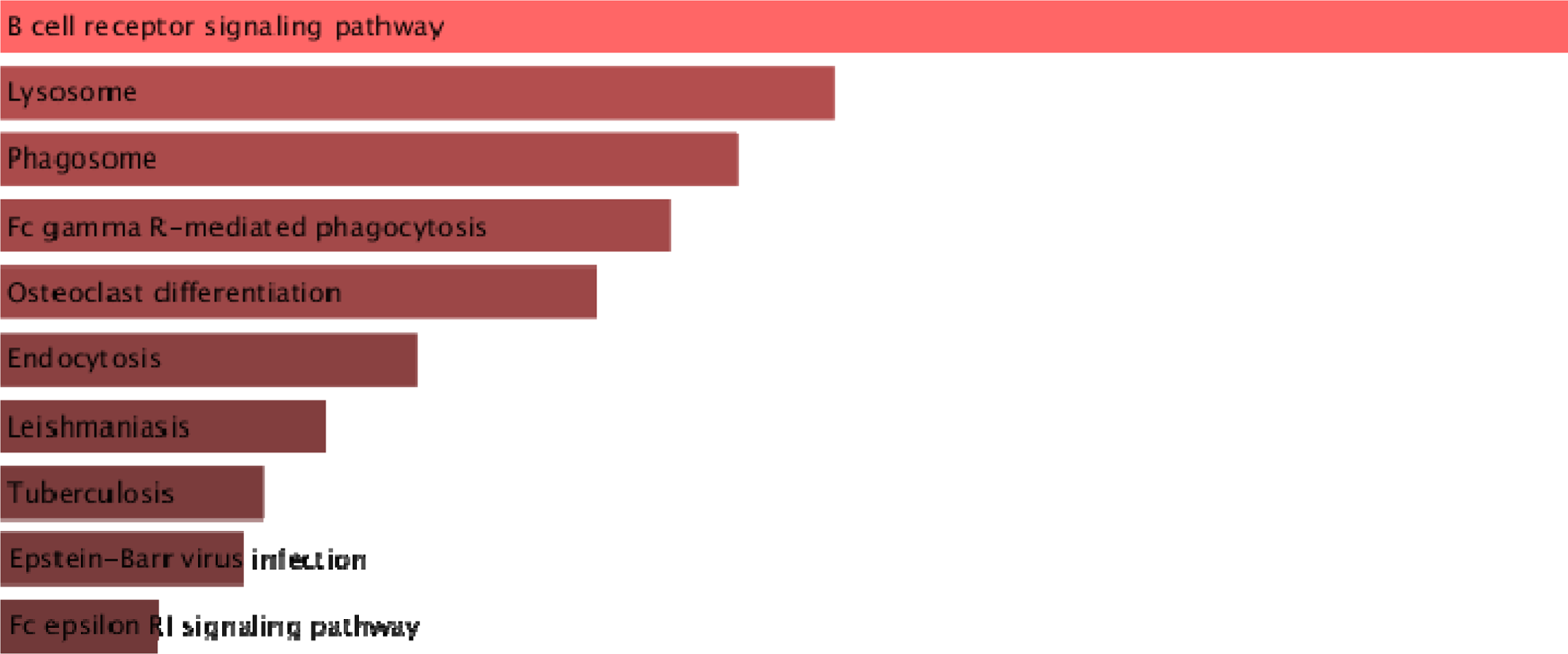
KEGG pathways *ENRICHR Gene Ontology and KEGG pathways analysis of significantly over-expressed genes of the African-American population.

### Significantly enriched KEGG pathways

Tuberculosis, Epstein-Barr virus infection, Influenza A, PD-L1 expression and PD-1 checkpoint pathway in cancer, B cell receptor signaling pathway, T cell receptor signaling pathways were observed to be significantly enriched in the upregulated genes, among other pathways (Figure. 5E) with multiple overlap genes (Table 3).

**Figure 5E.**
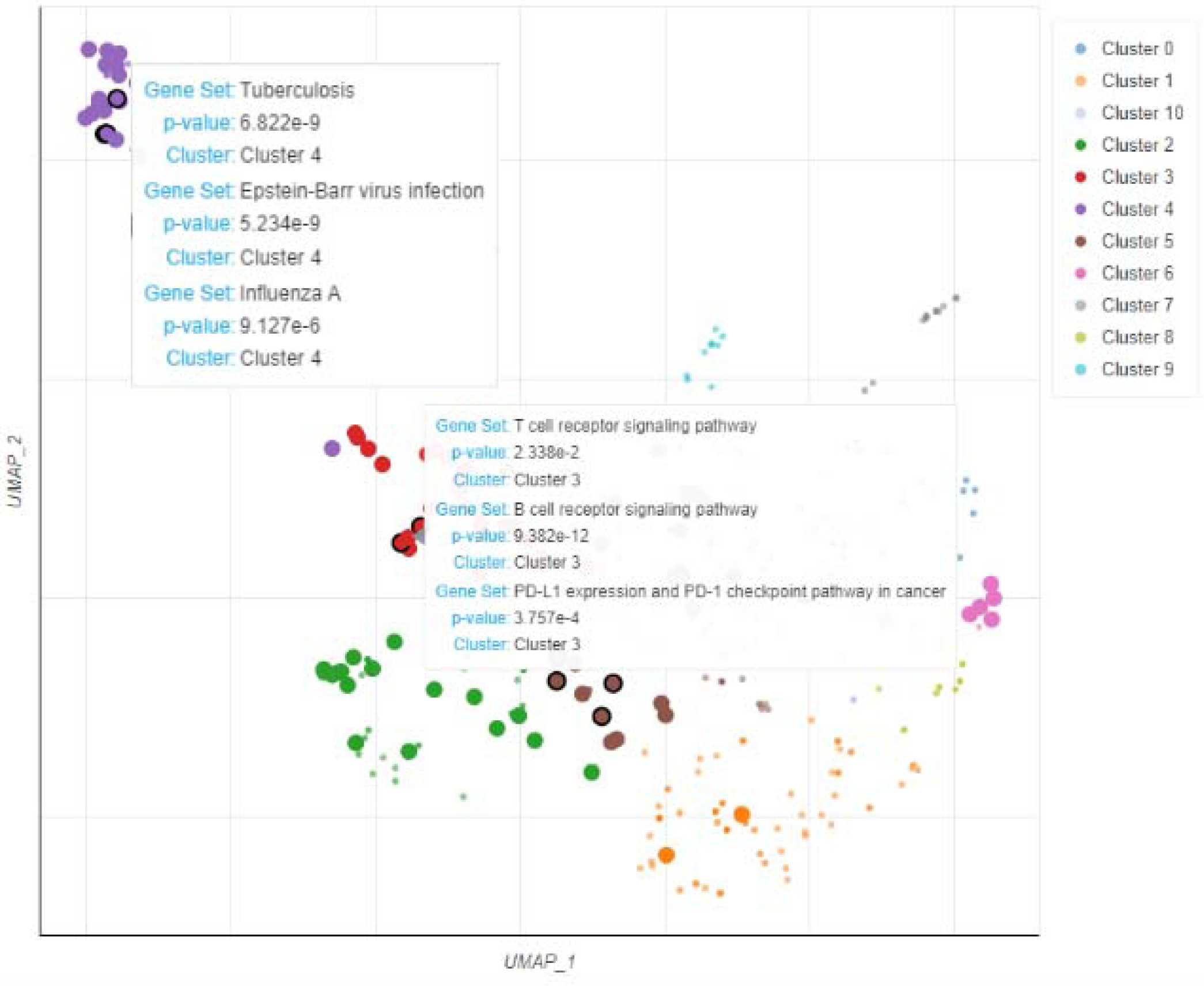
Scatterplot of all terms in the KEGG_2021_Human gene set library. Each point represents a term in the library.

**Table. 3.**
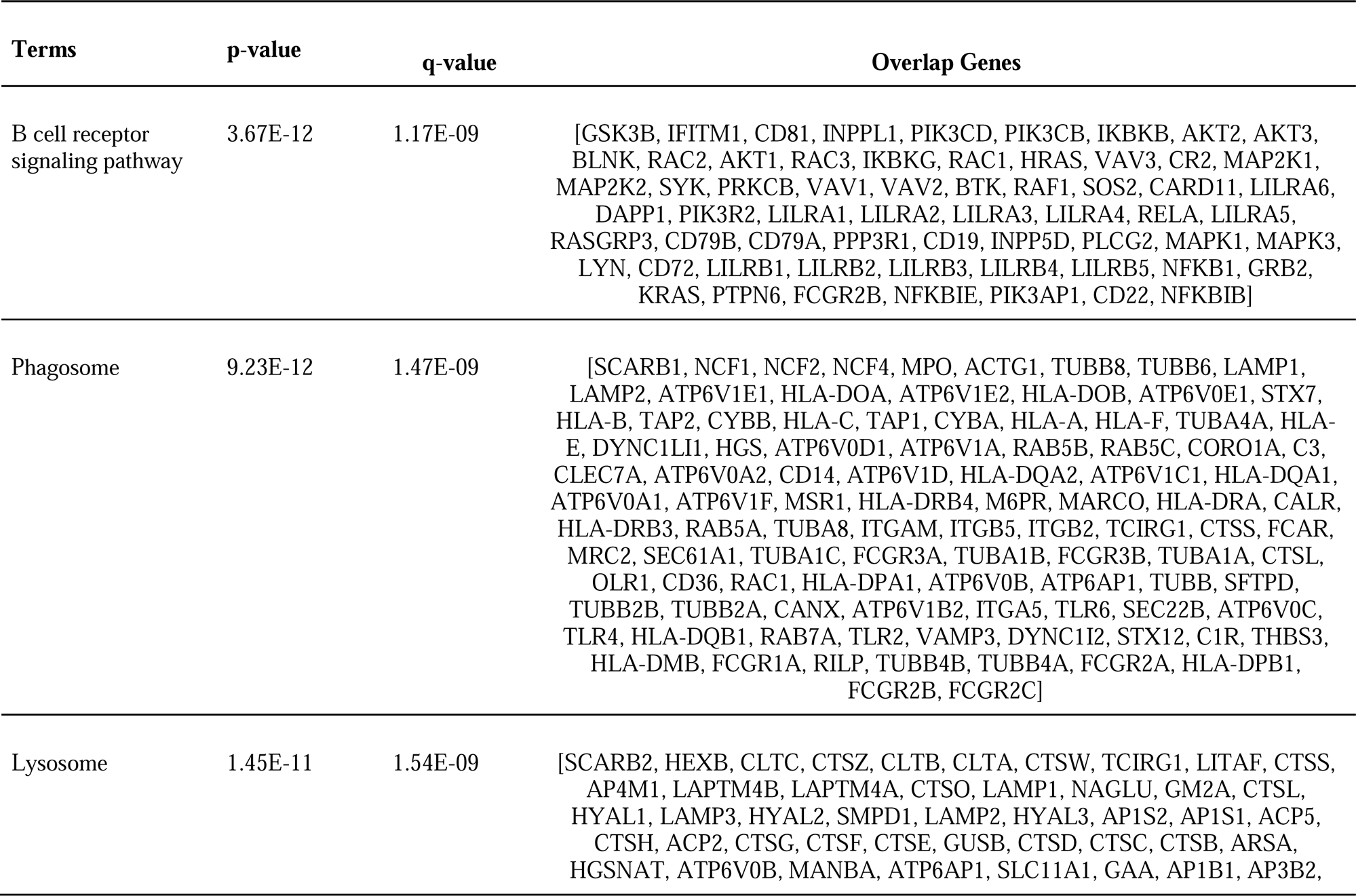
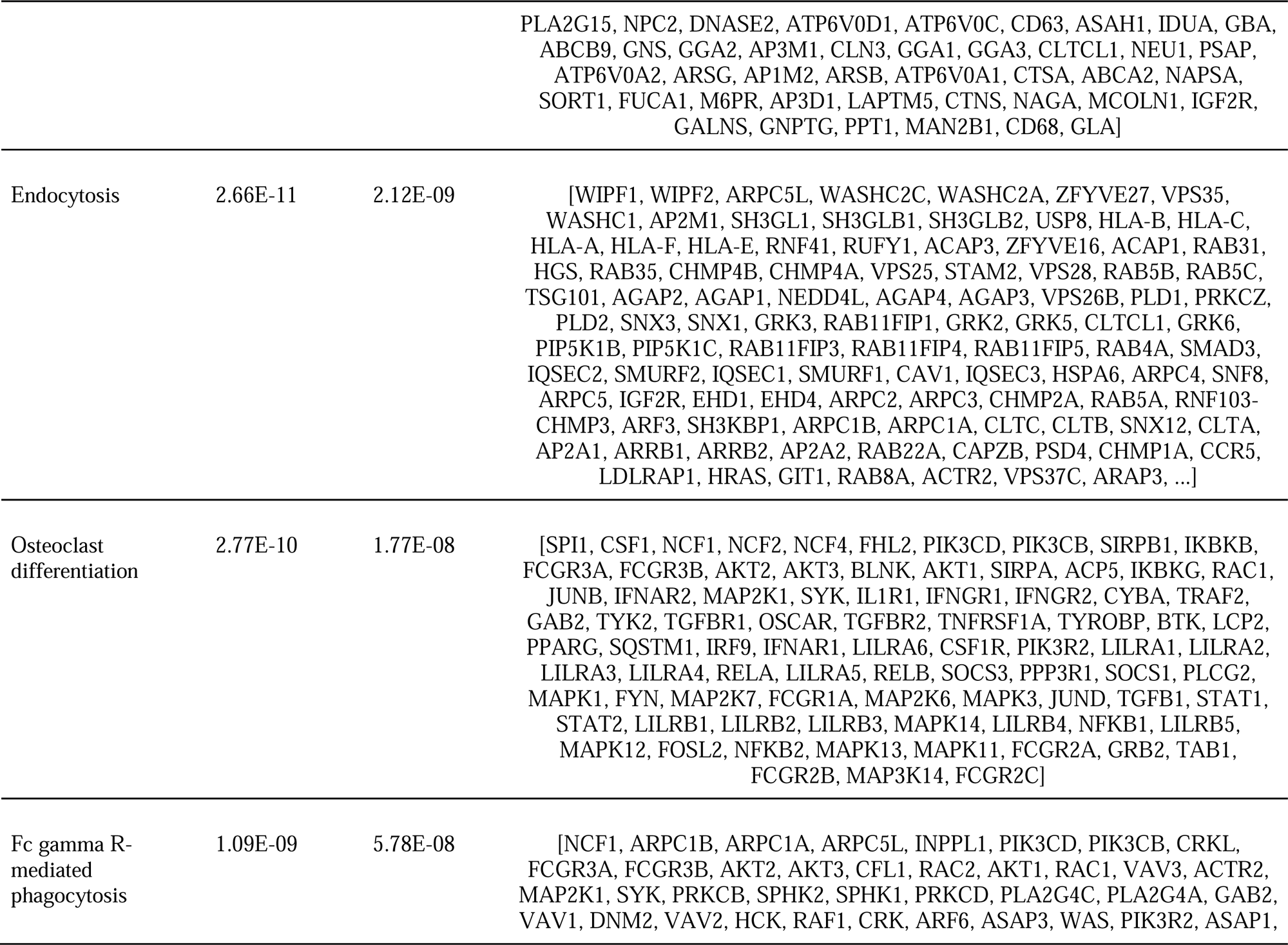
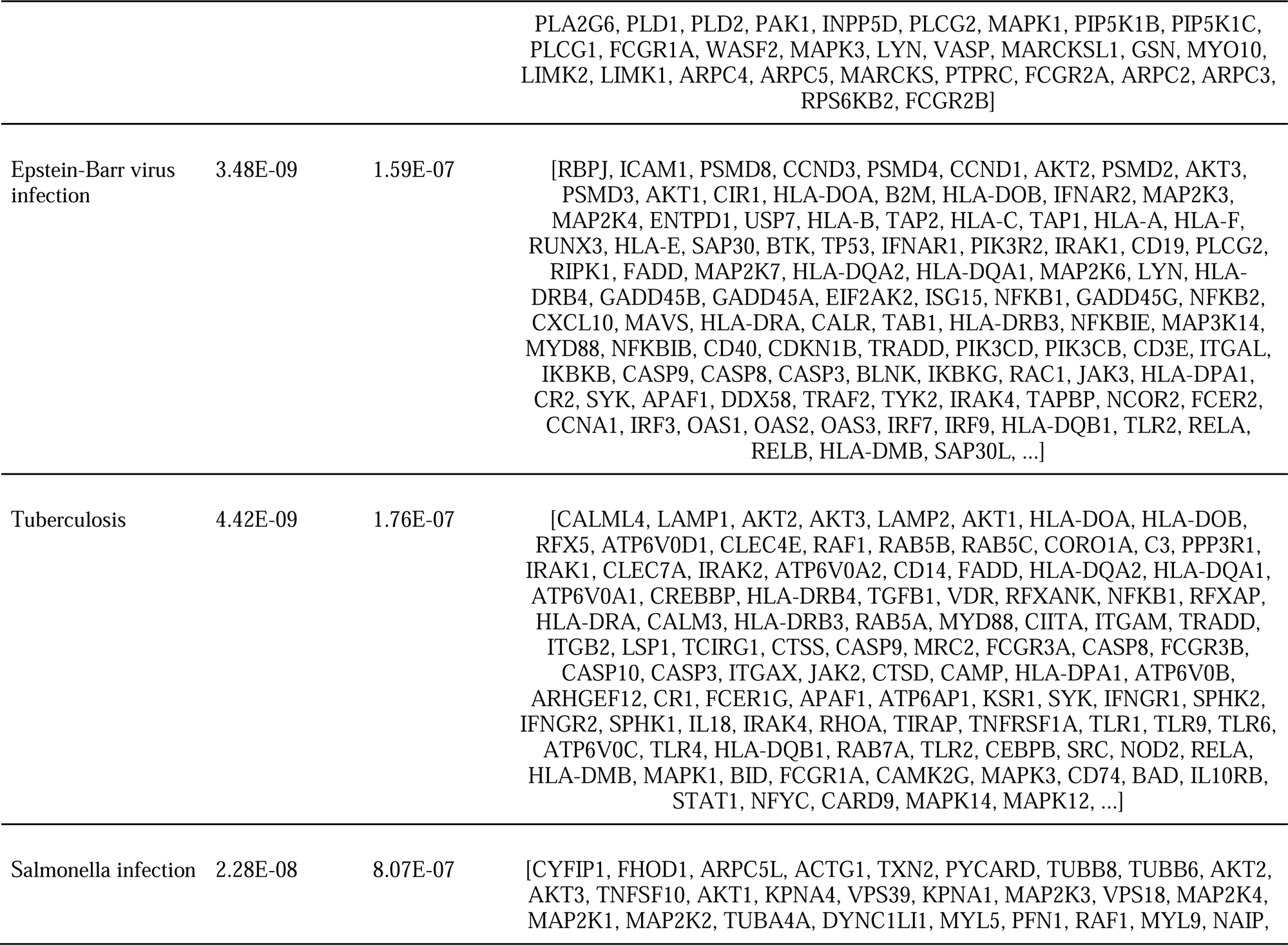
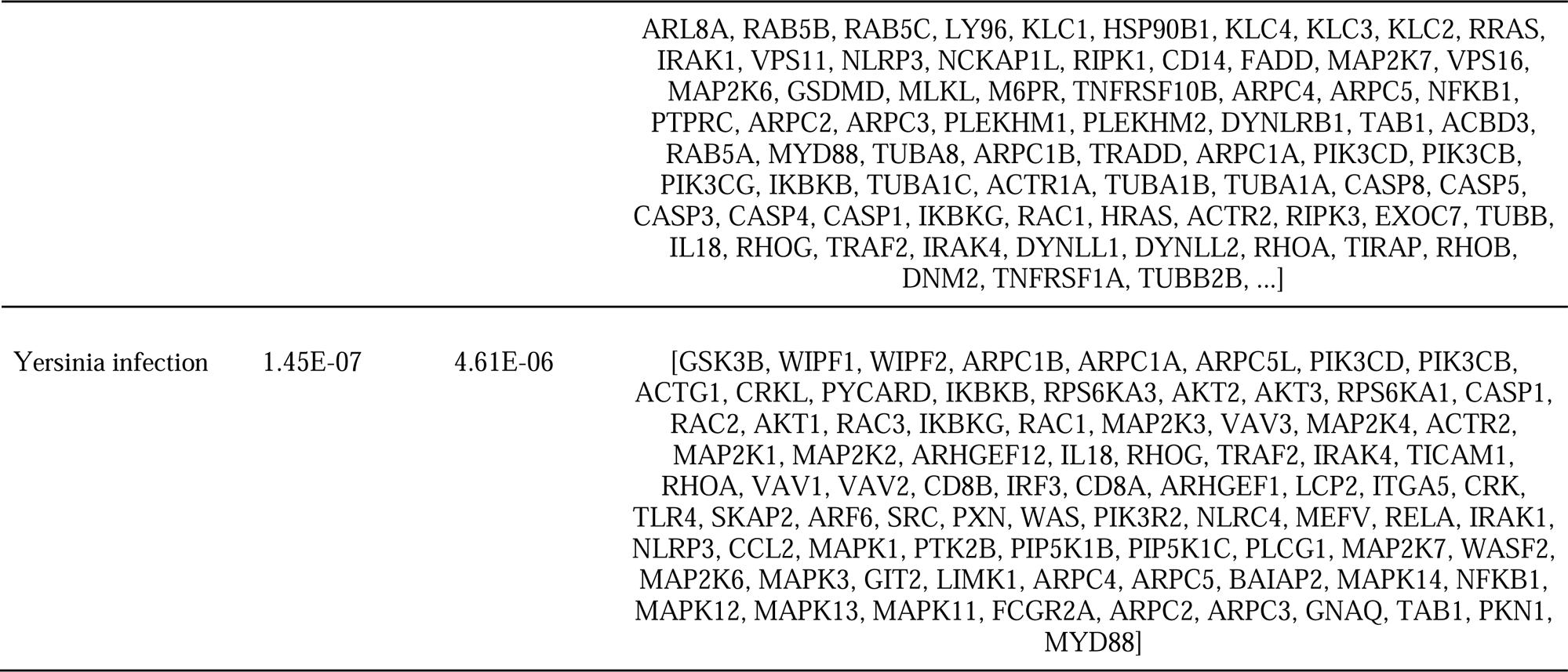
Top ten KEGG 2021 Human significant p-values and q-values, 2021.

### Functional analysis of significantly downregulated genes

To validate the molecular functions, biological processes, cellular components, and pathways linked with the differentially expressed genes, enrichment analysis was done on significantly under-expressed genes. According to Gene Ontology (GO) biological process, Gene Ontology (GO) molecular function, Gene Ontology (GO) cellular component, and KEGG pathway analysis based on p value, the fifteen most significant molecular functions, biological processes, cellular components, and ten most significant pathways associated with the differentially expressed genes are represented. (Figure 6A-6D).

**Figure 6A.**
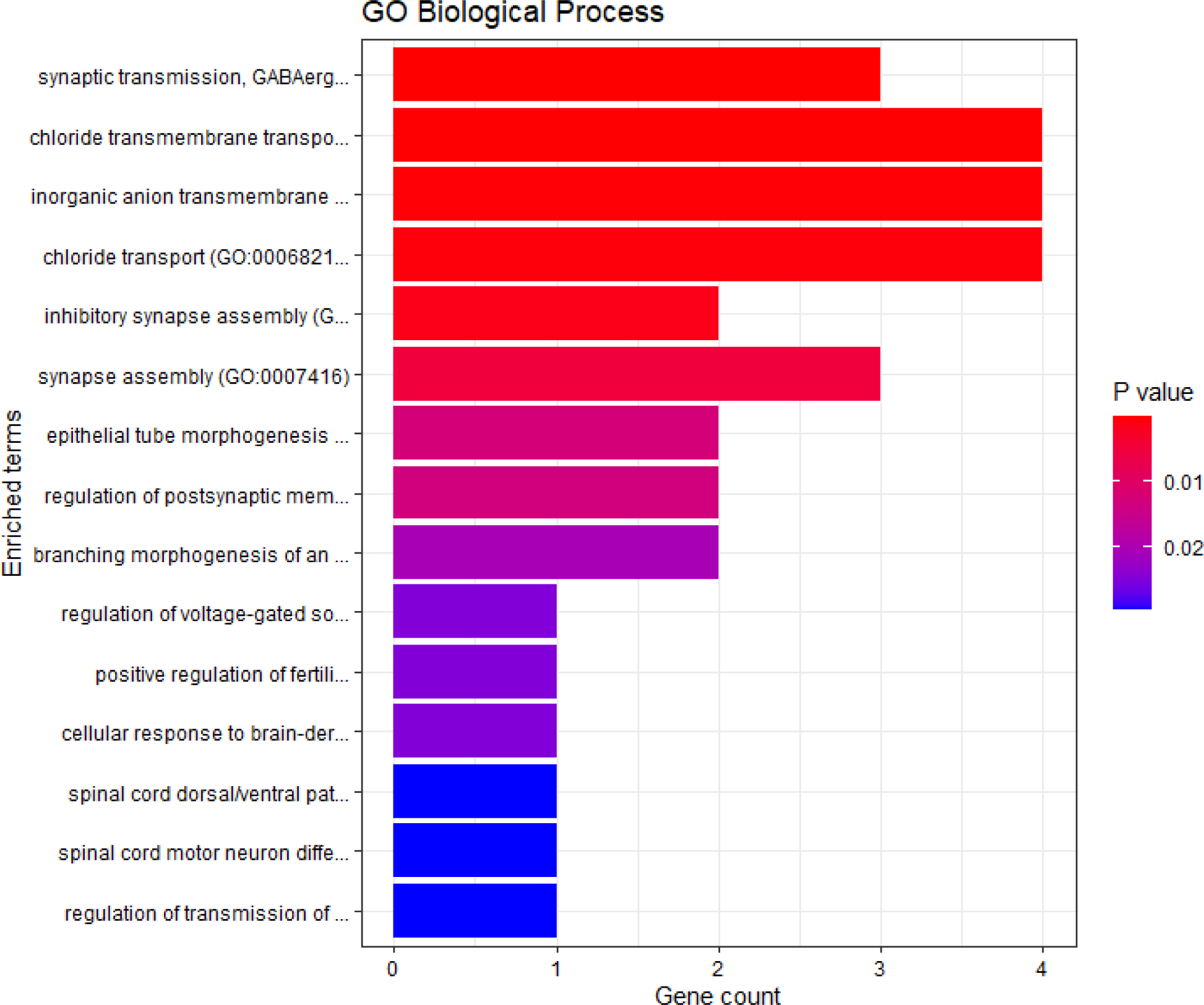
GO biological process

**Figure 6B.**
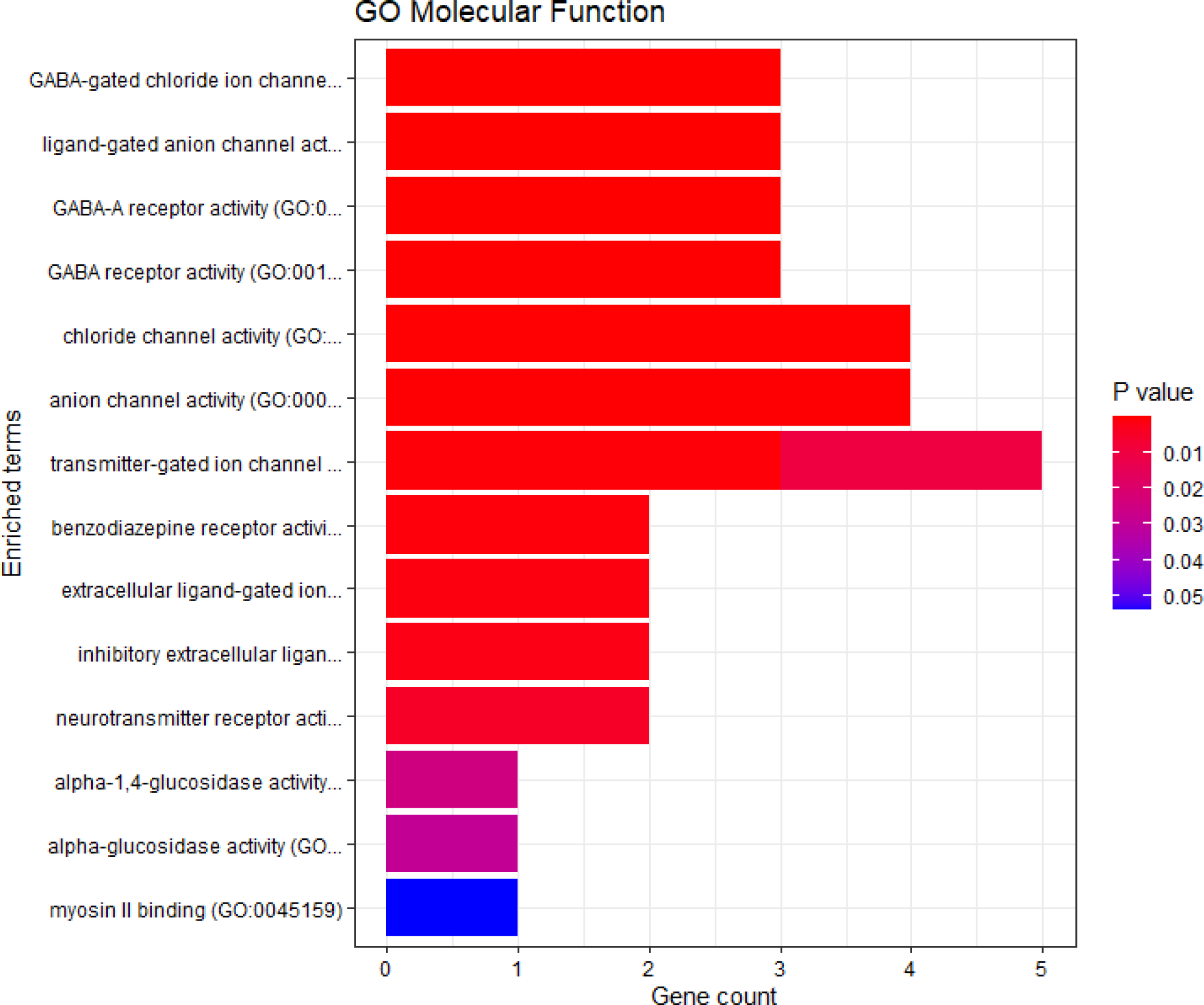
GO molecular function

**Figure 6C.**
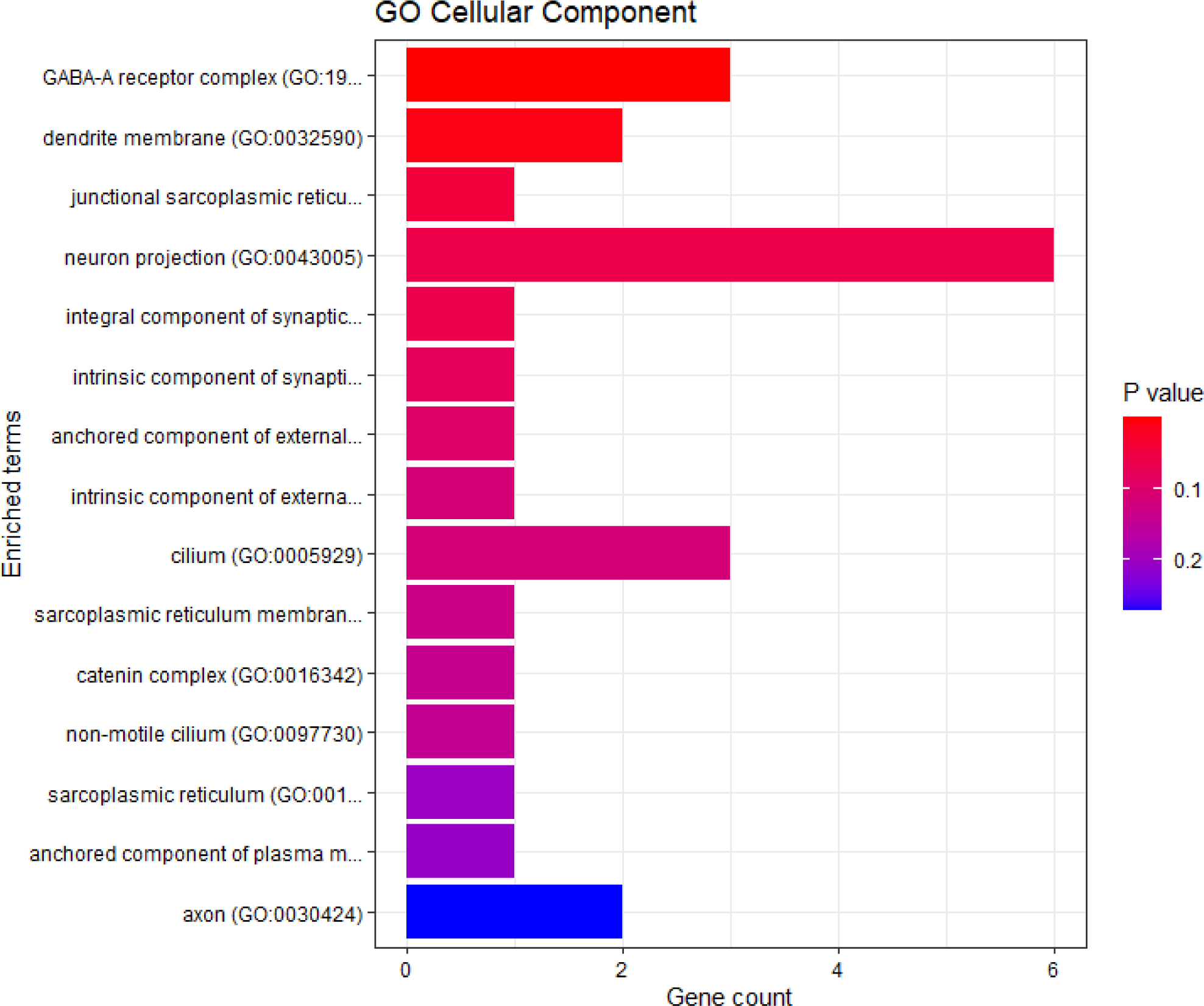
GO cellular component

**Figure 6D.**
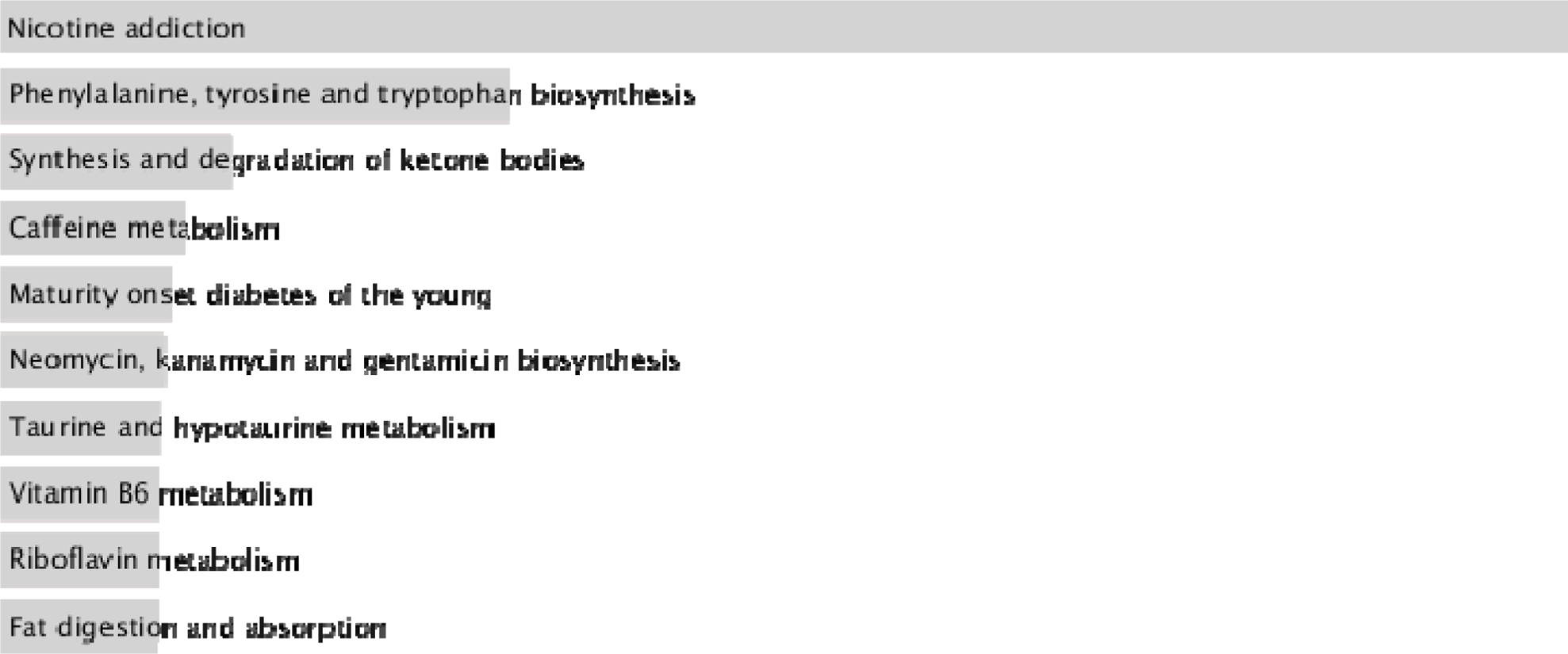
KEGG pathways *ENRICHR Gene Ontology and KEGG pathways analysis of significantly under-expressed genes of the non-African population.

### Significantly enriched KEGG pathways

Nicotine addiction, caffeine metabolism, prion disease, thermogenesis, Huntington disease, drug metabolism, T cell receptor signaling among other pathways were significantly enriched in the downregulated genes (Figure 6E).

**Figure 6E.**
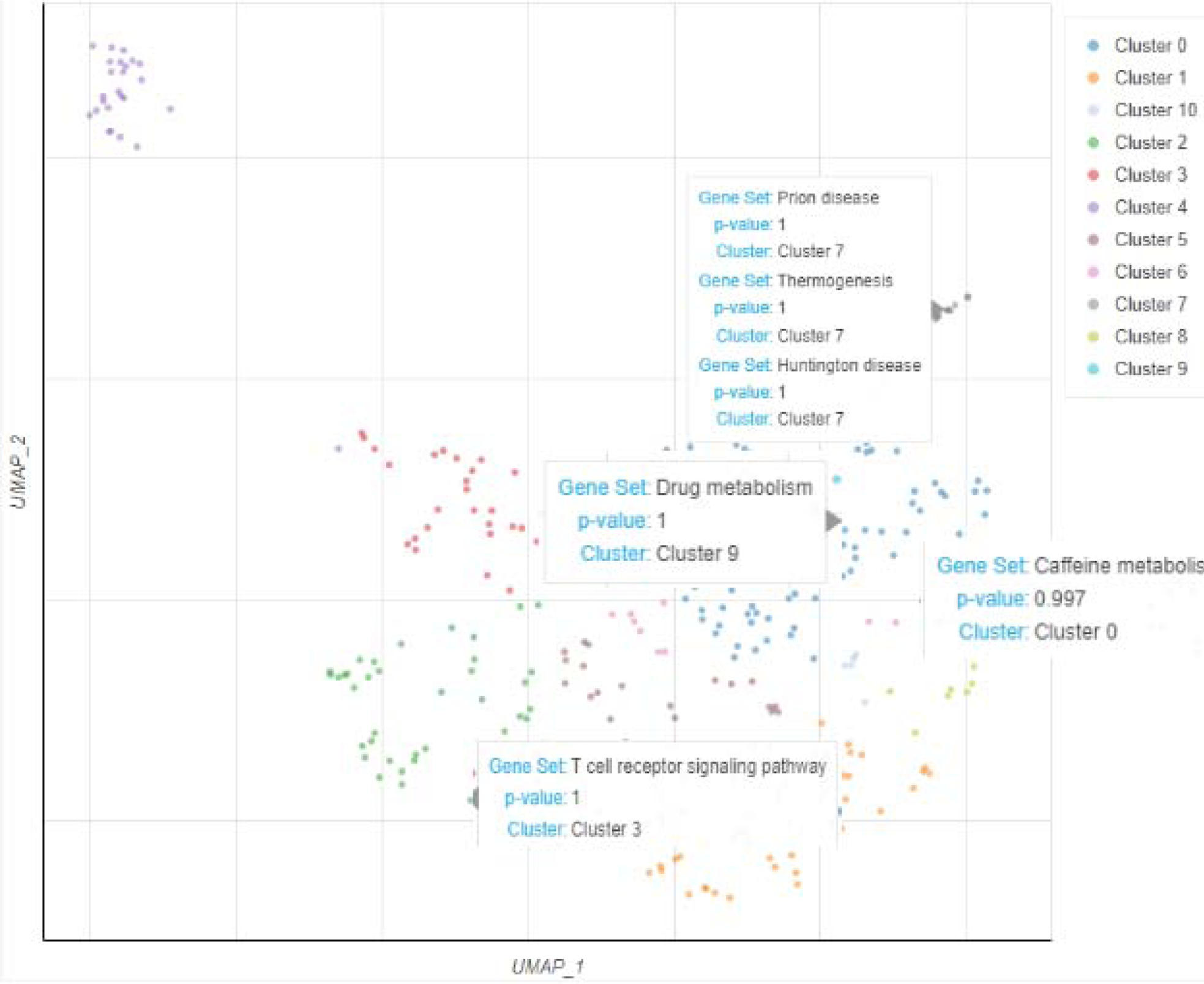
Scatterplot of all terms in the KEGG_2021_Human gene set library. Each point represents a term in the library.

## Discussion

The study set out to assess the difference in ACE2 expression in COVID-19 infected patients of African-American and non-African-American descent. Preliminary analysis between healthy African-American and COVID-19 infected African-American individuals did not show any difference in ACE2 expression. Likewise, a similar comparison between healthy and COVID-19 infected non-African-American individuals did not show any difference in ACE2 expression level. Our primary finding showed that the ACE2 gene was significantly downregulated in the African-American cohort compared to non-African-American cohort. The findings from this study might explain Yang and Cadegian’s [15], [53] observation of reduced COVID-19 severity with reduced ACE2 expression.

There are reports that COVID-19 has a higher infection rate and severity among the African-American population [34], [52]. Asides the probable genetic cause of the disproportionate COVID-19 rates and deaths in African-Americans, other social factors such as overcrowding, working conditions, access to healthcare and background chronic health conditions may have played a greater role.

We analyzed gene expression in the two populations and discovered that African-American patients had 7,718 differentially upregulated genes and 16,374 differentially downregulated genes when compared to non-African-American patients. Many of these genes may be important genes for COVID-19 research. Our study confirms that there are significant disparities in differentially expressed genes (DEG) expression levels between African-American and non-African-American COVID-19 patients.

Residue function and enrichment analysis of downregulated genes suggests that nicotine addiction, caffeine metabolism, thermogenesis, drug metabolism, T cell receptor signaling, prion and Huntington disease pathways are actively involved in viral disease manifestation of African-American COVID-19 population as compared to a non-African-American population. Normally residues of the former two pathways prevent disease severity by blocking ACE2 from binding to COVID-19 spike protein, decreasing viral virulence or replication and inflammation [29], [42]. Therefore, the suppression of their corresponding genes in the diseased population clearly supports their significance in close protection against COVID-19 infection. The abundant presence of xanthine and caffeine/nicotine which are directly involved in the caffeine/nicotine metabolism pathway also directly reduce viral propagation in African-Americans by inhibiting red blood cells-ACE2 complex that allows viral entry [43], [57]. Autoimmune genes associated with prion and Huntington disease pathways regulate subtle peripheral and Central Nervous System (CNS) inflammation caused by pro-inflammatory cytokines such as Interleukins (IL) and Tumor Necrosis Factor (TNF) – α hence slowing down neural degeneration [22]. Hypothetically, the viral attack of the patient’s nervous system by COVID-19 [55] might have significantly impacted these pathways resulting in downregulation of respective genes.

Thermogenic regulation is highly dependent on expression of ACE2 whereby its downregulation enhances microcirculation [47], thus subsequent downregulation of the thermogenic pathway in absence of any impairment. Cytochrome enzymes are active agents in toxin and drug degradation and their suppression by infection leads to compromised drug metabolism giving rise to treatment complications especially in COVID-19 patients [11], [53]. Even though genes for these enzymes are usually upregulated in African-Americans as compared to non-African-Americans [44] discrepancies observed here might be attributed to socio-environmental factors beyond our control during sampling. Negative regulation of other pathways such as T cell receptor signaling reduces viral cell entry as well as T cell production controlling immunopathology [56]. Evidently, the respiratory, inflammatory and infectious disease pathways play a role in COVID-19 progression.

Moreover, the expression levels of TGFB1, VASP and CYBA in African-American COVID-19 patients were all elevated in our study among other genes. TGFB1 and CYBA are associated with the osteoclast differentiation pathway; TGFB1 is involved in regulatory systems that support the balance of osteoblasts and osteoclasts [45] while over-expression of CYBA is linked to decreased blood stem cells [45] Moreover, the expression of TGFB1 in Tuberculosis disease is known to be elevated as it impacts the patients’ immunological response to the disease [36].

Meanwhile, the expression of VASP and MAPK1 have been reported to be associated with phagocytosis [27] and Fc Gamma R-mediated phagocytosis [24]. However, more study should be performed to confirm the function and character of VASP in with Fc Gamma R-mediated phagocytosis. Moreover, an upregulated gene linked to APOL1-associated kidney disease among African-Americans [48], [49], NLRP3 was found to be associated with salmonella infection in our study and its expression levels enhanced in our COVID-19 African-American patient transcriptomes. Our TGFB1, CYBA, and VASP results are consistent with those previously reported [24], so we speculate that these may be potential COVID-19 - related genes and more research is required.

### Conclusions and next steps

The expression of ACE2 gene is downregulated in the cohort of African-American COVID-19 patients in this study. This may explain the low severity of COVID-19 in this population compared to non-African-American populations. Future exploration of the ACE2 gene and the pathway residues of respiratory, inflammatory and infectious disease pathways might offer new chemical therapeutics for this virus.

## Declarations

### Availability and Requirements

Project name: Expression Level Analysis of ACE2 Receptor Gene in African-American and Non-African-American COVID-19 Patients

Project home page: https://github.com/omicscodeathon/ace2_covid

### Availability of data and materials

The dataset analyzed during the current study as a case study is publicly available at https://github.com/omicscodeathon/ace2_covid/blob/main/accessions. The data supporting the results reported in this manuscript is included within the article. The Project repository which also includes the entire code and other requirements can be downloaded from https://github.com/omicscodeathon/ace2_covid. The guidelines for this project and related updates, are available at: https://github.com/omicscodeathon/ace2_covid/blob/main/README.md

## Abbreviations

ACE2: angiotensin-Converting Enzyme 2
SARS-CoV-2: severe acute respiratory syndrome coronavirus 2
SRA: sequence read archive
Ang-I: angiotensin I
TGFB1: transforming growth factor beta-1
APOL1: apolipoprotein L1
NLRP3: NOD-, LRR- and pyrin domain-containing protein 3
STING1: Stimulator Of Interferon Response CGAMP Interactor 1

## Acknowledgements

The authors thank the National Institutes of Health (NIH) Office of Data Science Strategy (ODSS) and the National Center for Biotechnology Information (NCBI) for their immense support before and during the April 2022 Omics codeathon organized in collaboration with the African Society for Bioinformatics and Computational Biology (ASBCB). The authors acknowledge the assistance of Danny Lumian for editing the manuscript.

## Contributions

MN conceived the original idea and developed the pipeline, performed the bioinformatic analysis of the data, and wrote the original draft of the manuscript. AF supervised, wrote, reviewed and edited intensively the manuscript and performed administrative roles. KO performed the bioinformatic analysis of the data, reviewed the manuscript and provided critical feedback and helped update the github repo. BK performed the bioinformatic analysis of the data, wrote, reviewed and edited the manuscript. OD performed the bioinformatic analysis of the data. OIA drafted the original abstract and provided the needed resources and guidance for the successful completion of the project. OIA reviewed the manuscript and provided critical feedback. All authors read and approved the final manuscript.

## Ethics Declaration

### Ethics approval and consent to participate

Not applicable.

### Consent for publication

Not applicable.

